# Machine learning guided cell-free expression maps the biochemical landscape of carbonic anhydrase

**DOI:** 10.64898/2026.07.07.736810

**Authors:** John T. Lazar, Evan Komp, Irene Martinez, Kyle S. Zolkin, Pascal Notin, Samer Saleh, Grant M. Landwehr, Kyung G. Kim, Anru Tian, Benjamin A. Shapero, Ashty S. Karim, Debora S. Marks, Gregg T. Beckham, Michael C. Jewett

## Abstract

Carbonic anhydrases are among the fastest known biocatalysts, reversibly facilitating the hydration of CO_2_ to HCO_3_^-^ at rates up to 10^7^ s^-1^, which warrants their investigation for industrial carbon capture technologies. However, engineering carbonic anhydrases to maintain stability under harsh industrial process conditions remains a key challenge, and sequence-to-function datasets compatible with machine learning to inform forward engineering are lacking. Here, we developed a high-throughput platform that couples cell-free gene expression with a gaseous CO_2_ colorimetric assay to map the fitness landscapes of carbonic anhydrases. From 96 diverse natural homologs, we identified a robust variant from the *Aquificota* phylum and conducted an exhaustive mutational scan and functional assessment of this enzyme at 70°C and 90°C, covering >99% of all single-amino acid substitutions (totaling 4,365 mutations assayed in 39,285 reactions). This biochemical landscape was used to benchmark 22 zero-shot protein fitness models and identify critical mutations that improved enzyme stability at 90°C by more than three-fold. We then used both zero-shot protein language models and supervised learning to filter 419 model-generated variants from a ProteinMPNN library of 100,000 sequences, leading to a best-in-class enzyme that retained activity after incubation at 95°C. This work demonstrates that integrating cell-free enzyme engineering with machine learning enables opportunities for high-throughput experimental measurements to benchmark and improve protein language models, accelerate design loops, and expand functional exploration within protein families where experimental information is limited.

## Introduction

Carbonic anhydrases catalyze the hydration of CO_2_ into bicarbonate (HCO_3_^-^) at diffusion-limited rates^1–3^. This hydration reaction is a central component of natural mineral carbonation through rock weathering, a process in which bicarbonate reacts with various metal ion sources to precipitate permanent metal carbonates^4^, thereby removing ∼1 gigaton of CO_2_ per year from the atmosphere^5,6^. Given rising CO_2_ emissions that contribute to anthropogenic climate change^7^, carbonic anhydrases have been explored as a potential catalyst for large-scale CO₂ removal in both direct air capture (DAC) and flue gas sequestration. However, the industrial deployment of these enzymes has been hampered by the requirement for high-performance catalysis under elevated temperatures (>70°C) and alkaline pH typical of industrial carbon capture cycles^1–3^.

Numerous protein engineering campaigns have sought to improve the stability of carbonic anhydrase variants to bridge this gap. These strategies include bioinformatics searches for carbonic anhydrases from known thermostable species^8–10^ and rational site-directed mutagenesis^11–14^. Despite progress, past efforts have been biased towards a relatively small subset of the phylogenetic landscape. Furthermore, previous screens have often relied on proxy assays, such as hydration in CO_2_ saturated water^15^, esterase activity^16^, or supplying bicarbonate and measuring the reverse reaction^14^, which fail to accurately reflect the nuances of capturing gaseous CO_2_ as a substrate under industrial conditions. Thus, engineering a robust carbonic anhydrase capable of meeting the demands of industrial CO_2_ removal remains an outstanding challenge.

In recent years, enzyme engineering has been reshaped by machine learning (ML) approaches^17^, *de novo* design tools such as ProteinMPNN^18^ and RFDiffusion^19^, and protein language models (pLMs)^20^. In conjunction with off-the-shelf tools that can be used for function-agnostic tasks, building supervised models with labeled protein sequence-to-function data from high-throughput functional screens can improve optimizing a specific objective. However, a key challenge is measuring data of multiple dimensions (e.g., activity and thermostability) at a scale and speed that captures both successes and failures to inform forward engineering. To date, the lack of open-source, large-scale sequence-to-function mappings for carbonic anhydrases has limited the ability of deep learning architectures to expand beyond sequence space sampled by nature.

Here, we developed a high-throughput platform that combines cell-free gene expression (CFE) and a pH-based colorimetric assay to rapidly map various sequence-to-function landscapes of carbonic anhydrases (**Fig. 1A-C**). We used this platform to explore a broad range of natural carbonic anhydrases at elevated temperatures, identifying a highly thermostable cluster within the *Aquificota* phylum. We then performed an exhaustive deep mutational scan, testing 99.9% (4,365 of 4,370) of the single-amino acid mutants of the carbonic anhydrase from *Aquificota bacterium*, mapping the functional activity of >99% single amino acid substitutions across three different temperatures. We utilized this comprehensive dataset to provide mechanistic insights into the enzyme, identify key residues that confer significant improvements in stability, and benchmark various zero-shot fitness predictors. In parallel, we used deep learning sequence redesign via ProteinMPNN to combinatorially vary larger portions of the enzyme, surveying candidates further from the wild type in sequence space while minimizing inactive variants. By integrating iterative rounds of ML-guided selection and generative modeling, we identified redesigned variants that surpass best-in-class natural and engineered enzymes. Overall, our results indicate that changes to protein function are readily accessible through the combination of cell-free prototyping and deep learning sequence redesign.

**Figure 1:**
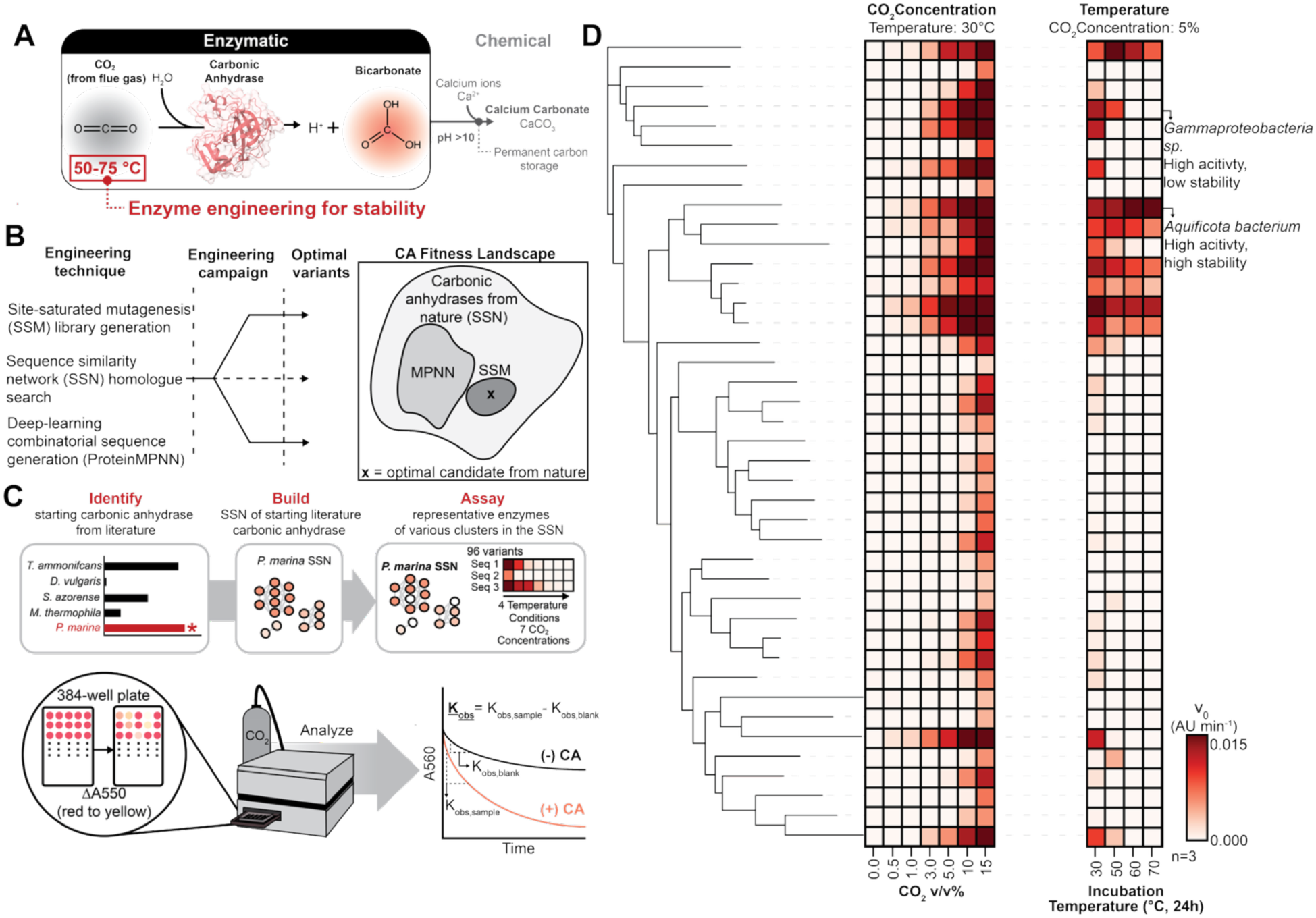
Exploration of carbonic anhydrase natural sequence diversity to identify optimal thermostable candidates. **(A)** Cartoon schematic of CO_2_ biomineralization via the enzymatic carbonic anhydrase (CA) hydration and chemical metal carbonate precipitation reactions. **(B)** Overview of carbonic anhydrase engineering campaign, outlining the types and timings of each engineering technique along with the broad feature space explored in each step. **(C)** Experimental pipeline of exploring natural sequence space. Thermostable variants from literature were identified and tested in our assay conditions to identify the optimal natural candidate (*P. marina* CA) for building a sequence similarity network. Representative members of each cluster in the SSN were then assayed in varying CO_2_ and post-temperature incubation conditions via our high-throughput assay, represented via cartoon. Initial reaction velocity was calculated through background subtracting the sample’s linear reaction velocity by the negative controls. **(D)** Heatmap v_0_ data of SSN carbonic anhydrases in conditions where (i) temperature was held constant at 30°C and CO_2_ concentration was varied between 0-15% v/v and (ii) CO_2_ concentration was held constant temperature incubation post-CFE was varied between 30-70°C for 24 hours. Shading of heatmap represents the average of n = 3 biologically independent replicates. Data in the main text represents only the 41 out of 96 carbonic anhydrases that demonstrated to have measurable CO_2_ activity in relevant CO_2_ conditions; data of all 96 variants can be found in supplemental **Figure S1**. Source data is provided in the source data file.

## Results

### Exploring the natural sequence space of carbonic anhydrases to identify an optimal engineering candidate

We began by surveying the literature for natural carbonic anhydrase sequence diversity to identify a robust starting point for engineering. Specifically, we selected 5 candidates spanning three distinct classes (alpha, beta, and gamma) reported to demonstrate activity at temperatures above 60°C. These enzymes were expressed in crude extract-based CFE reactions, and their activity was evaluated in both cell-free lysates and as purified enzymes. For functional screening, we employed a modified version of the Wilbur Anderson Assay^15,21^ that measures gaseous CO_2_ hydration in high-throughput (**Fig. S2**). The carbonic anhydrase from *Persephonella marina* significantly outperformed all other reported thermostable homologs tested (**Fig. S3**).

To systematically explore the natural sequence diversity surrounding the *P. marina* carbonic anhydrase, we identified the 1,000 most similar UniProt sequences by BLAST (BLOSUM62, gap penalty score = 11, e-value = 10^-5^) and constructed a sequence similarity network (SSN) using the Enzyme Similarity Tool from the Enzyme Function Initiative^22^ (67% similarity cutoff; **Fig. S4**). From this SSN, we selected 96 carbonic anhydrases covering all major subnetworks, including representative sequences belonging to the *P. marina* subnetwork, for cloning as linear templates and expression via cell-free synthesis. Following expression, we measured CO_2_ hydration directly in cell-free lysate across 7 CO_2_ concentrations and 4 pre-incubation conditions (**Fig 1D, Figs. S1, S4**). Of these 96 enzymes, 41 retained measurable activity at ≤15% v/v CO₂, and 7 of these remained active when measured after a 24 h incubation at ∼70°C. Embedding all sequences using Evolutionary Scale Modeling-2 (ESM-2) showed that the thermostable variants cluster tightly in latent space, whereas only four cluster together in the SSN (**Figs. S4-5**). The best-performing natural enzyme, isolated from an *Aquificota* species, exhibited an *in vitro* >2.8-fold improvement in activity over the *P. marina* carbonic anhydrase after 6 h at 70°C (**Fig. S6**). We selected this *Aquificota* carbonic anhydrase (AQ-WT) as a model sequence to enhance thermostability.

### Site-saturation mutagenesis of an entire thermostable carbonic anhydrase

To begin our enzyme engineering campaign on AQ-WT, we explored the single mutant space of the enzyme holistically to identify beneficial mutations, understand mutational tolerance, and generate large datasets for integration with protein language model benchmarking. To this end, we constructed a full site-saturation mutagenesis (SSM) library of AQ-WT covering all 230 residues (**Fig. S7**). The library comprised 4,371 unique variants (due to wild type repetition) and was verified by testing for DNA amplification via Quantiflor (**Fig. S8**) and full sequencing of ∼2% of all clones.

We assessed the activity of every carbonic anhydrase variant under three conditions: (i) no pre-incubation, (ii) pre-incubation for 120 h at 70°C, and (iii) pre-incubation for 120 h at 90°C, yielding 13,113 sequence-to-function measurements (41,400 data points including all replicates and controls) (**Fig. 2A, Fig. S9**). Biological replicate concordance was high for the unincubated and 90°C datasets (Pearson’s correlation coefficient R > 0.8; **Fig. S9**), but lower at 70°C (R ∼ 0.6; **Fig. S10**); we therefore focused our analysis on the two higher-quality datasets. Strikingly, AQ-WT proved highly mutable, with ∼95% of single substitutions retaining ≥80% of WT activity without incubation (**Fig. 2A-2B**). Deleterious mutations were primarily localized to the active-site region (**Fig. 2C, Fig. S11**). These results were consistent with evolutionary analyses of carbonic anhydrases from hydrothermal vents showing conservation of catalytic residues with sequence variability in solvent-exposed regions.^23^

**Figure 2:**
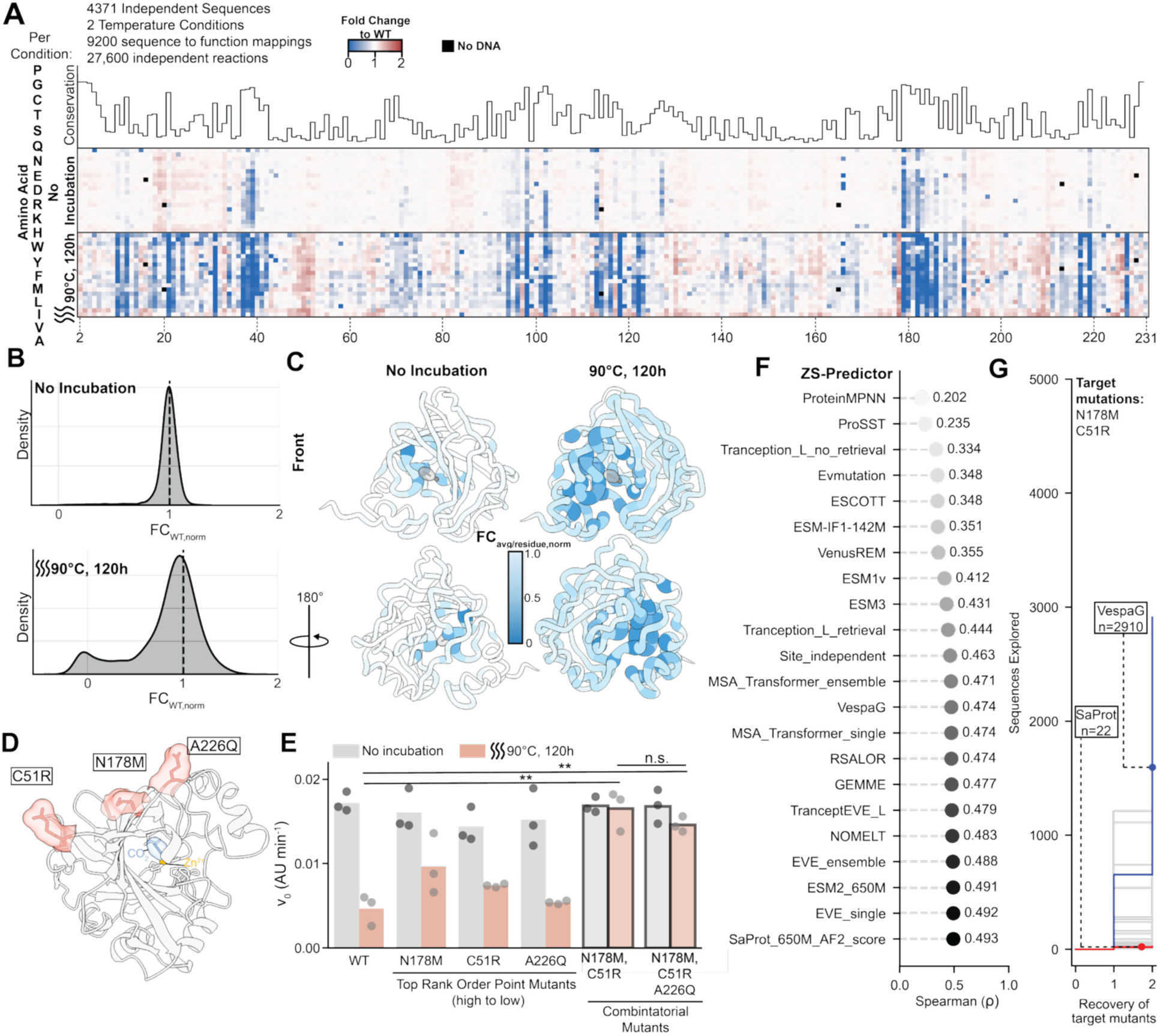
Whole-enzyme site saturated mutagenesis of *Aquificota bacterium* carbonic anhydrase. **(A)** Per residue conservation paired with site-saturated mutagenesis heatmaps of all residues in AQ-WT in both a no incubation condition and post 120h incubation at 90°C. Crossed out boxes represent no DNA present/excluded data. Color of the heatmap box represents the mean of n =3 biologically independent replicates. **(B)** Kernel density distributions of all fold-change data in both incubation conditions. **(C)** Worm plots where averaged fold change per residue, normalized to WT (FC_avg/residue, norm_) is plotted onto the color and thickness of the worm at each residue. **(D)** Chai-1 predicted structure of AQ-WT with CO_2_ and Zn^2+^ cofactor bound in the active site. The top 3-point mutations colored in red. **(E)** Bar plot of top mutants along with combinations of the top mutations. Each bar represents the mean of n = 3 biological replicates. **(F)** Spearman correlation of 22 Zero-shot predictor scores against the observed activity in the 90°C, 120h incubation condition. **(G)** Ability of the same scorers as in **F** to recover the top 2 mutations observed in 90°C, 120h incubation condition, where the y-axis is the number of rank ordered mutations according to predictor score and x-axis is the recovery of C51R and N178M mutations. Source data provided in source data file.

After the 90°C incubation, ∼32% of substitutions reduced activity below 80% of WT activity (**Fig. 2A**, **Fig. 2C**). Destabilizing mutations again mapped near catalytic residues (**Fig. 2C**). We identified multiple stabilizing substitutions: N178M, C51R, and A226Q improved activity by 1.7-, 1.5-, and 1.6-fold, respectively. Mapping these variants onto the predicted structure (**Fig 2D**) revealed that they are located adjacent to the catalytic site and near a disulfide bond at the interface of the modeled tetrameric assembly (**Fig. S11**). Individual validation of these variants confirmed improved enzyme stability for N178M and C51R relative to AQ-WT (∼2-fold and ∼1.6-fold post 90°C, 120h incubation), whereas A226Q did not provide a statistically significant improvement (p > 0.05, two-tailed student’s T-test, **Fig. 2E**). Combining stabilizing substitutions revealed minimal dependence on A226Q; the N178M/C51R double mutant performed best, achieving 3-fold improvement in measured enzyme activity following the 120-h, 90°C incubation, which almost fully restored wild-type activity levels (**Fig. 2E**).

### Benchmarking protein language models against an experimentally complete fitness landscape

pLMs have recently been shown to excel at enzyme functional annotation^24,25^; yet, their ability to predict user-specified, quantitative performance metrics remains poorly understood due to a scarcity of comprehensive experimental datasets. It is additionally difficult to know *a priori* which, if any, modeling methods might correlate with the objective function of interest (e.g., enzymatic activity with a target substrate, catalytic rate, thermal stability, etc.). To this end, we leveraged our carbonic anhydrase deep mutational scan to assess the predictive accuracy of multiple zero-shot scoring methods. Across the complete mutational landscape and 20 zero-shot fitness predictors, we found modest but distinct correlations (Spearman’s rank correlation coefficient, ρ, up to 0.49; **Fig. 2F**). Notably, SaProt recovered the top 2 confirmed stabilizing mutations (N178M and C51R) within the top 22 predictions of the 4,371-variant library (**Fig. 2G**). Only two additional models (ESM-1v, n = 152; and ESM2, n = 350) were able to recover the top 2 stabilizing mutations among the top 10% of their rank order (**Fig. 2G, Fig. S12)**. ProSST, which has performed well in the broader ProteinGym zero-shot benchmark^26^ demonstrated poor predictability (ρ = 0.235, Top 2 recovery n = 2493). Structure-aware predictors (i.e., SaProt, ESM-IF, ESM3) tended to perform best across both metrics, suggesting that stability determinants are largely structural and exhibit limited epistasis in this enzyme. However, ProteinMPNN, which is also structure-aware, performed poorly, indicating that architectural differences among models strongly influence zero-shot behavior.

Notably, the Spearman rank correlation coefficient poorly predicted a model’s ability to recover the top stabilizing mutations (**Fig. S13**). We therefore evaluated multiple complementary metrics, including the area under the receiver operating characteristic curve (AUROC), the normalized discounted cumulative gain (NDCG), and position-averaged scores (**Fig. S14–19**, **Table S1**). Predictive power varied across distinct structural domains: mutational effects on stability were most predictable for loop and helix residues near the active site, whereas they were least predictable for β-sheet positions. Collectively, our results suggest that zero-shot scoring can substantially reduce the experimental burden associated with engineering the stability of similarly mutable ⍺-carbonic anhydrases.

### MPNN-guided design of a non-thermostable carbonic anhydrase

Considering that our complete site-saturation mutagenesis (SSM) dataset of AQ-WT demonstrated that the enzyme possesses high mutational tolerance, we hypothesized that carbonic anhydrases are excellent targets for combinatorial redesign. To survey this higher-dimensional mutational space, deep learning inverse-folding models such as ProteinMPNN have been shown to improve both the stability and activity of various proteins by introducing broad combinatorial sequence variations (>30% mutational frequency)^18,27,28^. Thus, we next leveraged these inverse-folding architectures to design highly stable carbonic anhydrase variants.

As a testbed for ProteinMPNN design, we first selected a non-thermostable *Gammaproteobacteria* carbonic anhydrase (Gb-CA). We chose Gb-CA because of its proximity to AQ-WT in our sequence similarity search (**Fig. 1C**) despite its poor stability at 70°C (**Fig. 3A**), providing a larger dynamic range for optimization. To begin, we generated 60 computationally designed variants across a range of ProteinMPNN hyperparameters, varying the sampling temperatures (T = 0.1–1.0) and fixed-residue counts (30–70 amino acids, corresponding to 13-31% of the most highly conserved sites across the gene sequence, defined using EVCouplings conservation^29^). Five sequences were generated per combination of sampling temperature and fixed residues (**Fig. 3B**). All 60 sequences were predicted to fold well using AlphaFold 2 (pLDDT (Predicted Local Distance Difference Test) >90, pAE (Predicted Aligned Error) <5; **Fig. S20**). Experimental evaluation showed that 31 designs were active without incubation and 11 remained active after 24 h at 70°C (**Fig. 3C**). Higher sampling temperatures (T = 0.5–1.0) led to widespread inactivation, consistent with excessive diversification.

**Figure 3:**
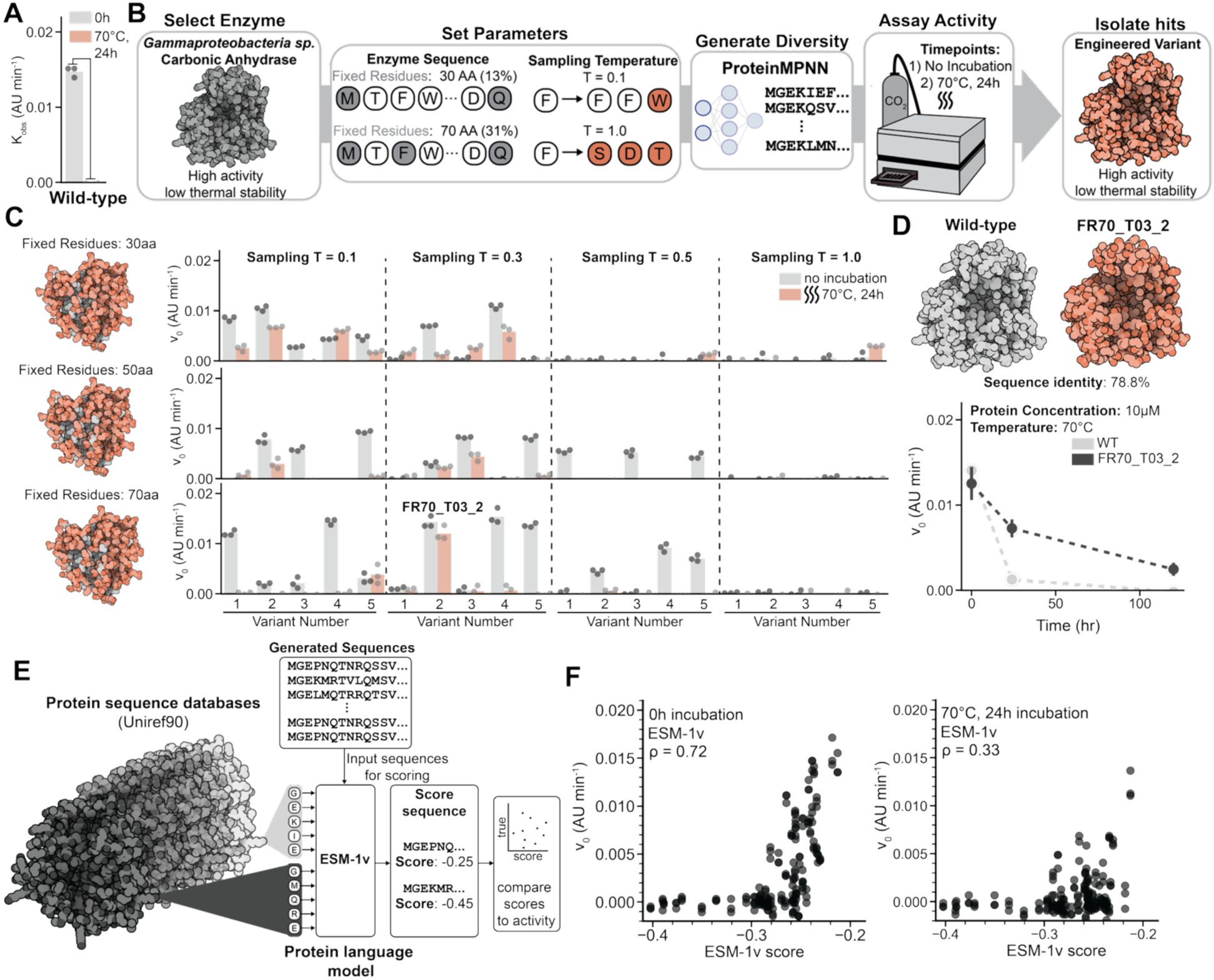
ProteinMPNN redesign of a non-thermostable carbonic anhydrase to improve thermostability. **(A)** Reaction velocity of *Gammaproteobacteria sp.* carbonic anhydrase pre-incubation and post-24h, 70°C incubation. Bars represent the mean of 3 biological replicates. **(B)** Cartoon schematic of ProteinMPNN design pipeline optimization. **(C)** Experimental data on combinatorial parameter optimization of sampling temperature and number of fixed residues. Activity was measured for both no incubation and post-70°C, 24h incubation where bar plots represent the mean of n = 3 biological replicates. **(D)** AF3 predicted structures of both WT *Gammaproteobaceria sp.* Carbonic anhydrase and FR70_T03_2 along with time course data of 10 µM purified stocks of the respective enzymes. Error bars represent the standard deviation of n = 3 biological replicates. **(E)** Cartoon schematic for benchmarking ESM-1v protein language against carbonic anhydrase activity data. Experimentally tested sequences are inputted into ESM-1v for scoring, which is then compared to experimental (pre and post incubation) data. **(F)** Parity plots of true activity vs ESM-1v score in both no incubation and 24-hour incubation condition. Spearman’s correlation coefficients are reported on the plots. Source data is provided in the source data file.

The most thermostable design (FR70_T03_2; generated at T = 0.3, 70 fixed residues) shared 79% identity with Gb-CA, showed increased soluble expression in CFE via ^14^C-leucine radiolabeling (**Fig. S21**), and remained active after 24 h and 120 h at 70°C (**Fig. 3D**), whereas the wild-type sequence exhibited no detectable activity. Our results demonstrate that ProteinMPNN enables substantial stability improvements when applied to non-thermostable carbonic anhydrase scaffolds.

We next explored the ability of zero-shot pLMs to identify the active and stable generated designs. Specifically, the sequence-only pLM ESM-1v (using the wild-type marginal score distribution) was used to score the 60 Gb-CA ProteinMPNN designs. ESM-1v was chosen because it has demonstrated an ability to threshold enzyme designs for the likelihood of activity in past studies^30,31^. Indeed, we observed that ESM-1v scores exhibited strong correlation with activities of the ProteinMPNN designs (Spearman’s ρ = 0.72 without pre-incubation; **Fig. 3F**) and moderate correlation post-incubation (ρ = 0.33; **Fig. 3G**), consistent with its performance on the SSM landscape (**Fig. 2**). The sequence-only CARP-640M pLM^32^ yielded similar predictability (**Fig. S22**). Taken together, our results support the emerging consensus that pLMs have predictive strength for both single point mutations and large generative libraries for functional activity^30^; however, identifying *a priori* which systems and mutational loads will yield strong correlations remains a key challenge.

### ProteinMPNN redesign of AQ-WT guided by supervised learning

We were next interested in combining our sequence-to-function mappings with ProteinMPNN to improve carbonic anhydrase activity, and we therefore applied an iterative design and validate pipeline to AQ-WT (**Fig. 4A**). Using the optimized parameters from the Gb-CA engineering campaign (T = 0.3, 70 fixed residues), we generated 100,000 ProteinMPNN sequences exhibiting an average sequence similarity of approximately 77% relative to AQ-WT (**Fig. 4B**, **Fig. S23**). Following sequence generation, we filtered this library using ESM-1v zero-shot prediction scores and chose the top 192 sequences for experimental testing (**Fig. 4C**). Two variants could not be successfully cloned and were excluded from subsequent analysis. Of the 190 variants tested, 98.4% (or 187) were active and 15 exhibited statistically significantly increased stability after 120 h at 70°C pre-incubation relative to AQ-WT (p < 0.05, two-tailed student’s T-test, **Fig. 4D**). However, ESM-1v scores correlated only weakly with stability (ρ = 0.13) at this elevated temperature.

**Figure 4:**
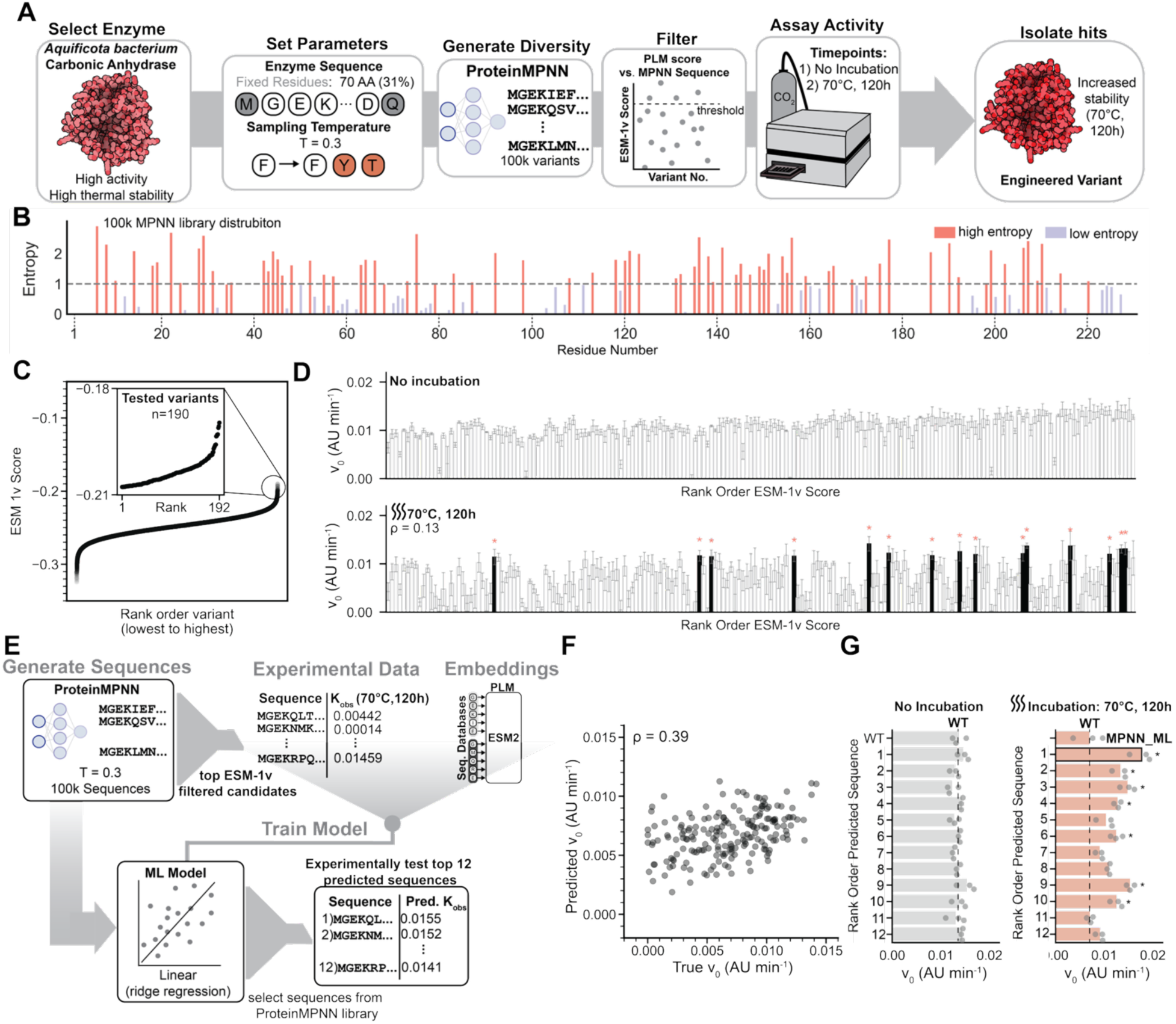
ML-guided ProteinMPNN engineering of the thermostable AQ-WT carbonic anhyhdrase. **(A)** Cartoon schematic detailing updated ProteinMPNN pipeline for improving stability of AQ-WT. Adjustments include utilizing optimal design parameters from previous engineering campaign, ProteinMPNN sequence generation significantly more *in silico* variants (100,000) and the addition of a ESM-1v filter. **(B)** Position independent Shannon entropy for AQ-WT ProteinMPNN generated sequences. Low entropy represents low residue variance (few, biased mutations to WT residue) while high entropy represents increases in greater site variance and mutation rate. **(C)** Protein language model scoring the entire ProteinMPNN library with ESM-1v. Top 192 variants (specified in subplot) were ordered to be cloned and expressed in CFE, and 190 of the 192 were able to then be experimentally tested. **(D)** Experimental data of the top-ranked ESM-1v variants in both a no incubation and post-120h, 70°C incubation. In the 70°C, 120h incubation plot, black bars with an “*” above denote variants with statistically significant (p > 0.05, two-tailed student’s T-test) increases in stability compared to AQ-WT. Error bars represent the standard deviation of n = 3 biological replicates. **(E)** Cartoon representation of supervised machine learning model trained on experimental data from top ESM-1v candidates used to filter next round variants. **(F)** Parity plot of 70°C, 120h experimental data from **D** compared to predicted activity of the supervised linear model. Spearman’s correlation coefficient reported on the plot. **(G)** No incubation and 70°C, 120h data of the supervised model’s top 12 predictions from the ProteinMPNN library. Dotted line represents the mean reaction velocity of AQ-WT in each condition. “*” above denote variants with statistically significant (p > 0.05, two-tailed student’s T-test) increases in stability compared to AQ-WT. Bars represent the mean of n = 3 biological replicates. Source data is provided in the source data file.

To better identify highly stable variants from ProteinMPNN, we trained supervised learning models on the experimental annotations from the 190 sequences tested in the prior round (**Fig. 4E**), exploring both sequence embedding strategies (ESM2 at various sizes and pooling strategies) and predictor architectures. Training linear ridge regression models markedly improved predictive accuracy of carbonic anhydrase stability (ρ = 0.39; **Fig. 4F**) compared to ESM-1v scoring alone. Non-linear models did not further improve predictive power (**Fig. S24**). The linear model was then used to rank the remaining untested variants of the library of 100,000 ProteinMPNN sequences for experimental selection. Testing the top 12 predictions yielded 7 variants with improved stability compared to AQ-WT (**Fig. 4G**), surpassing the initial ESM-1v-only hit rate (7.8%). The best variant, “MPNN_ML,” showed the highest stability for any ProteinMPNN-designed sequence after 120 h at 70°C (0.0177 ± 0.0021 AU min⁻¹).

To reduce sampling bias introduced by ESM-ranking in the first round, we examined the embedding distribution across the full 100,000-member library. Experimentally tested variants occupied a narrow region of ESM2 latent space (**Fig. S25**). To broaden coverage, we selected 200 additional carbonic anhydrase variants using D-optimality as a target and measured their activities. D-optimality selects variants that maximize the determinant of the information matrix, resulting in diverse sequences that improve a model’s understanding of the design space. Incorporating these data and training on all 390 sequences raised performance of the linear model to ρ = 0.62 and the nonlinear model to ρ = 0.59 (**Fig. S25**). All top 12 predictions from both models exceeded AQ-WT stability with several variants matching the stability of MPNN_ML (**Fig. S26**). We attempted to incorporate the SSM data into model training by representing all data as fold change to wild type control, but found that mixing the datasets decreased performance compared to training on only combinatorial ProteinMPNN variants (**Fig. S27**); all correlations were calculated in 5-fold cross validation on all available data at the time of their testing, e.g., the data from previous rounds.

We next tested whether these models could generalize to higher-entropy generative ProteinMPNN libraries. Using increased sampling temperatures (T = 0.5, 0.7, 1.0), we generated 3 computational libraries of 500,000 variants each (**Fig S28**). Model agreement decreased as diversity increased; yet both models successfully identified stabilized variants even at T = 1.0, corresponding to near-random combinatorial sampling (**Fig. S29**).

### Biophysical and kinetic characterization of top engineered carbonic anhydrase variants

Following our enzyme engineering prototyping efforts in the cell-free platform, we purified AQ-WT, the SSM double mutant N178M_C51R, the best ML-designed variant MPNN_ML, and the literature benchmark DvCAv10 for detailed biophysical comparison (**Fig. 5A**). DvCAv10 is an engineered beta carbonic anhydrase variant obtained after 10 rounds of directed evolution, reported to retain stability at 107°C for 1 hour^14^. Size-exclusion chromatography (SEC) showed that AQ-WT forms heterogeneous oligomers, whereas DvCAv10 forms a single homogeneous species (**Fig. 5B**). The N178M_C51R mutant retained multiple oligomeric states; however, MPNN_ML displayed a single, soluble oligomeric species closely resembling DvCAv10 in sizing (**Fig. 5B**). Circular dichroism revealed high structural stability for MPNN_ML with a melting temperature exceeding 95°C (T_m_ > 95 °C; **Fig. S30**).

**Figure 5:**
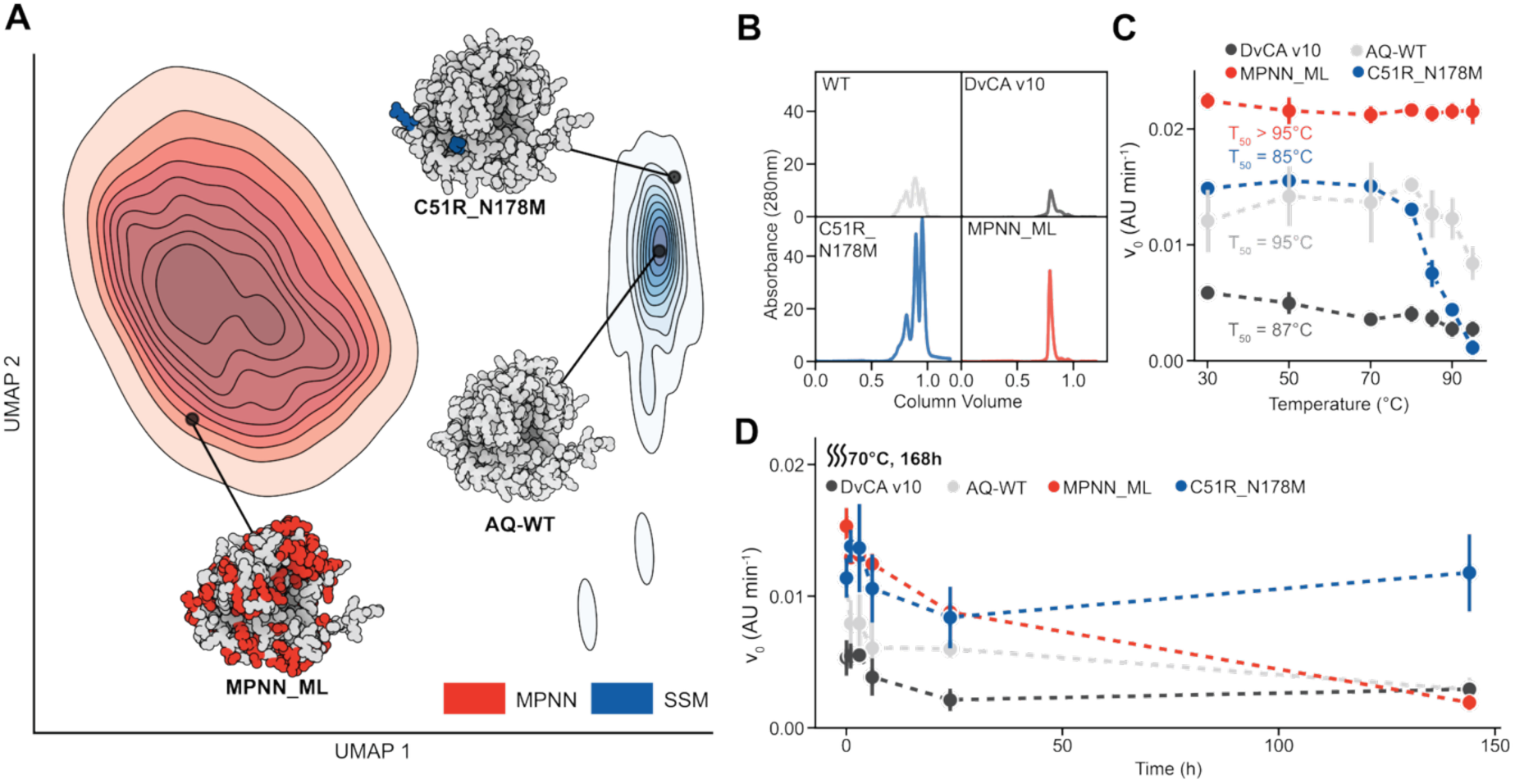
Structural and enzymatic stability characterization of optimal engineered variants. **(A)** UMAP of the ESM2^20^ embeddings of all sequences from the SSM (n = 4372) and ProteinMPNN (n = 100,000) libraries. Locations of variants AQ-WT, C51R_N178M, and MPNN_ML are all specified in the latent space. AF3 predicted structures of each of these respective variants, colored by mutation (grey = same as AQ-WT, red = mutation from ProteinMPNN engineering campaign, blue = mutation made from SSM engineering campaign) are provided on the UMAP. **(B)** Size exclusion chromatography curves of each characterized carbonic anhydrase variant. For each curve, 100 µL of 1 mg/mL protein was loaded onto the column. **(C)** T_50_ enzyme characterization of selected variants. Enzymes were incubated at the specified temperatures for 30 minutes before reaction velocities were measured. Error bars represent the standard deviation of n = 3 biological replicates. **(D)** Long-term stability measurements of specified enzyme variants at 70°C. Variants were incubated in Eppendorf Thermomixers for the specified time before removed to have their reaction velocities measured. Error bars represent the standard deviation of n = 3 biological replicates. All enzymes in **C** and **D** were incubated at 10 µM, and the enzyme concentration in the reaction was 1 µM. Source data is provided in the source data file.

Kinetic stability is a critical metric for determining industrial feasibility of carbonic anhydrase variants. We therefore generated T_50_ curves to identify the temperatures resulting in a 50% loss of carbonic anhydrase activity post 30-minute temperature incubation (**Fig. 5C**). Kinetic stability ranked MPNN_ML highest (T₅₀ > 95 °C), followed by AQ-WT (95 °C), DvCAv10 (86 °C), and N178M_C51R (85 °C) (**Fig. 5C**). Week-long assays of the purified proteins at 70°C revealed a different ordering: N178M_C51R retained 103% activity and outperformed the state-of-the-art DvCAv10 by ∼4-fold (**Fig. 5D**). Longer lysate-based assays (2–3 weeks at 70–90°C) further highlighted the superior long-term stability of N178M_C51R over MPNN_ML, AQ-WT, and DvCAv10 (**Fig. S31**).

## Discussion

To leverage modern data-driven advancements for enzyme engineering, comprehensive datasets are needed to train models. Such datasets have been scarce, especially for carbonic anhydrases. As a step toward addressing this gap, we established a high-throughput cell-free platform to map the carbonic anhydrase biochemical landscape and train predictive workflows. We demonstrated the utility of this framework by studying natural and computationally designed sequences to execute an accelerated, iterative protein engineering campaign. In total, we explored the sequence-function landscape of 4,987 unique carbonic anhydrase sequences across 4 different incubation conditions through more than 45,000 total reactions. Our work had several key features.

First, we comprehensively characterized all possible single point mutants of a thermostable, 230 amino acid carbonic anhydrase. These data represent one of the very few whole-enzyme SSM libraries where enzymatic rates have been determined, particularly at multiple temperatures.^33–36^ Notably, these data allowed for rigorous assessment of zero-shot pLM predictions across a complete mutational landscape with experimental rates under varying thermal stress, providing a new benchmark to existing efforts^15,37^. We found that while pLMs such as ESM-1v and CARP-640M reliably identify broadly active variants, they fail to capture fine-grained thermostability trends. This insight underscores the need for high-quality experimental data to train supervised models and iteratively improve predictions.

Second, by combining this high-throughput dataset with deep learning-guided sequence redesign via ProteinMPNN, we identified both local point mutations and global combinatorial variants with markedly improved thermostability. Our site-saturation mutagenesis library revealed that AQ-WT is highly mutable outside the catalytic core, enabling identification of stabilizing mutations such as C51R and N178M, which together dramatically improved long-term thermostability. The stability increase in C51R_N178M is intriguing, as the mutation C51R removes four predicted intermolecular disulfide bonds among the tetramers. However, mutation of the two other cysteines in AQ-WT thought to form intramolecular disulfide bonds removes all activity post-incubation, suggesting that the intramolecular disulfide bonds contribute significantly more to carbonic anhydrase thermostability. This is a phenomenon reported before for *Thermovibrio ammonificans* carbonic anhydrase, AQ-WT’s closest relative^38^. Meanwhile, the combinatorial ProteinMPNN redesign variant MPNN_ML selected from our 100,000-member computational library via our supervised linear model has 70 mutations compared to AQ-WT yet achieved short-term structural and enzymatic stability, though long-term stability was surpassed by the site-saturation mutagenesis-derived mutant. These results demonstrate that point mutation and deep learning-guided redesign approaches can complement each other, offering the potential to map and exploit larger fitness landscapes when integrated.

Third, enhanced thermostability positions these engineered carbonic anhydrases as a further step towards industrial CO₂ capture applications. Future work will need to validate operational robustness under industrial conditions^4^ or assess integration with downstream bicarbonate utilization pathways^39–41^ to produce value-added metabolites^42–44^.

Overall, our work demonstrates that systematic, large-scale experimental mapping of sequence-to-function landscapes provides both a critical benchmark for machine learning models and a practical path for engineering enzymes with enhanced stability. By addressing this data scarcity, we demonstrate how the integration of high-throughput experimentation with deep learning-guided design can accelerate the discovery and rational optimization of industrially relevant enzymes, a paradigm that is readily extendable to diverse protein families beyond carbonic anhydrases.

## Methods

### DNA cloning and cell-free protein expression

Amino acid sequences for all carbonic anhydrase variants generated and assayed are included in the source data. For carbonic anhydrase homologs, ProteinMPNN designs, and combinatorial SSM variants, DNA gene fragments encoding the coding sequences along with 40bp of homology to pJL1 backbone on both the 5’ and 3’ ends were ordered through Twist Biosciences. Variants included a N-terminal CSL tag (CAT-Strep-Linker fusion containing Strep-tag II)^45^ for downstream protein purification. All high-throughput liquid handling was performed utilizing the Viaflo liquid handling robot equipped with either a 12.5µL or 50µL 384-well head. Gene fragments were diluted to 10 ng/µL. pJL1 was amplified via PCR using primers ACH86 and ACH87 as specified in the **Table S1**. PCR product of the backbone was then run on a DNA gel, and the correctly sized band was purified via gel extraction (Zymo Gel DNA Recovery Kit). 3 µL Gibson Assembly reactions^46^ were performed with 1µL of gene fragment and 0.4 µL of 50 ng/µL of purified, amplified pJL1 backbone and incubated at 50°C for 1h. Following, Gibson Assembly reactions were then diluted 1:10 in nuclease-free water. 2.5 µL of the diluted Gibson Assembly product is then added into 25 µL PCR amplifications along with 0.5µM of the primers ACH113/ACH114 (**Table S1**) to generate a linear expression template (LET) product for downstream cell-free expression.

For generation of the whole enzyme site-saturation mutagenesis library of the *Aquificota sp.* carbonic anhydrase, a cartoon schematic of the workflow can be found in **Fig S7**. Primer design, PCR amplification of the WT plasmid, dpn1 digestion, one-part Gibson Assembly, and LET amplification all directly follow the protocol outlined by Landwehr *et al.*^47^ Following amplification, 1 µL of LET was diluted 1:10 in nuclease-free water. Quantiflor fluorescence detection of amplified DNA was then performed by pipetting 27 µL of master mix (1x TE buffer, 1x Quantiflor dye, Promega) with 3µL of diluted LETs in clear 384-well plates (VWR). Plates were vortexed for 20 seconds and then incubated in darkness for 15 minutes before measuring fluorescence (504nm excitation, 531nm emission) on a plate reader (Biotek Synergy H1). Any samples that threshold below 50% of the max fluorescence across the plate were 1) ran on a DNA gel and 2) submitted for sangar sequencing to confirm absence of LET amplification. In addition, 8-16 randomly selected LETs per plate were selected for sangar sequencing to confirm the proper mutations were introduced across the subset. If any problems were observed across the plate (high WT background signal, incorrect mutation, etc.), the plate was discarded, and the Gibson assemblies were re-performed.

For screening protein variants, cell-free protein expression was performed as previously described, and on the scale of 3.75 µL reactions (with 0.5 µL of LET DNA) utilizing the PANOx-SP formulation^48^. Reactions were performed in 384-well plates and were incubated at 30°C for 20 hours before samples were either prepared to be assayed or heat challenged.

### Recombinant Protein Purification

Recombinant proteins were expressed in two separate scales. For small scale purifications (100-200 µL of ∼0.5-2 mg/mL of purified protein), 500 µL CFE reactions were prepared in FlowerPlates (Beckman Coulter) and incubated at 30°C, 500 RPM overnight for approximately 16 hours. To purify, the Strep-Tactin XT 4Flow high capacity Spin Column Kit (IBA Life Sciences) was utilized, following the instructions provided. Purified samples were then buffer exchanged into their storage buffer (90 mM Tris-HCl, 150 mM NaCl pH=8) utilizing 10kDa weight cutoff filters. SDS-PAGE gels and Bradford Assays were then utilized to confirm size, purity, and concentrations of the samples.

### Carbonic Anhydrase Activity Assay

Pre-activity assay, enzymatic variants for screening were expressed in 3.75 µL cell-free protein expression reactions as previously described in the methods and then either immediately assayed or subjected to the specified heat challenge in a convection oven (VWR) before assaying. For purified variants, unless otherwise specified, 50 µL of 10 µM purified enzyme were aliquoted into 1.5 mL micro centrifuge tubes were prepared and then either used for no incubation activity measurements or incubated at the specified temperature in Eppendorf Thermomixers.

A modified version of the Wilbur Anderson Assay^15,21^ has been utilized to assay carbonic anhydrase activity from specified gaseous CO_2_ volumes in high-throughput (Fig S1). For lysate-based assaying, pre- or post incubation crude lysate is mixed with 50 µL of reaction buffer (90 mM Tris-HCl, XX µM phenol red, pH = 8), and 30 µL of the mixed volume are transferred to black 384-well clear bottom reaction plate (Greiner). In the case of purified protein, reactions were prepared directly in the reaction plates, mixing 3 µL of 10 µM protein stocks with 27 µL of reaction buffer (1µM enzyme in reaction). Reaction plates were then spun down and inserted in Neo2 Plate Reader outfitted with a CO_2_ gas control module for 5 to 15 minutes. Unless otherwise specified, CO_2_ concentrations were set to 5% v/v, reaction temperatures were 30°C, and the plate absorbance was measured at 550nm. Reaction velocities of samples were then calculated from the linear slope of the initial reaction (typically within the first 5 minutes). Reported initial reaction velocity was then calculated by subtracting the sample value by the blank (no DNA for CFE reaction or no protein for *in vitro* study) reaction velocity.

### ProteinMPNN Sequence Library Generation, PLM Embedding/Filtering, and Structural Modeling

To generate ProteinMPNN guided libraries, ColabDesign (https://github.com/sokrypton/ColabDesign) was utilized on GoogleColab with a T4 GPU. Wild-type (WT) amino acid sequence, Alphafold 3^49^ monomeric predictions of WT structure, specified sampling temperatures, fixed residues, and desired amount of sequences were all inputted as specified in the manuscript/supplementary information. MPNN generated sequences were exported as a comma separated value (.csv) file for downstream processing. Fixed residues were selected based on positional conservation frequencies via multiple sequence alignment (MSA) from the EVCouplings pipeline^29^.

Zero-shot protein language model filtering for the initial screen of ProteinMPNN candidates was performed utilizing Colab notebooks developed by Johnson *et al*.^30^(https://github.com/seanrjohnson/protein_scoring). ProteinMPNN designed sequences were fed into the sequence-only protein language models ESM-1v^31^ and CARP-640M^32^ with a single-pass. All zero-shot predictions were performed utilizing a T4 GPU on GoogleColab. Predicted protein structures of various carbonic anhydrases both engineered and from nature were generated utilizing AlphaFold3^49^ or Chai-1^50^ as noted throughout the manuscript.

For the retroactive benchmarking of zero-shot fitness predictors in **Fig. 2G-H**, the ProteinGym pipeline was run on CA-WT including the following sequence, evolutionary, and structure models: ProteinMPNN, EVMutation, ProSST, ESM-1v, VenusREM, ESM-IF, Site independent, Tranception with and without retrieval augmentation, EVE with and without stochastic MSA subset training as an ensemble, MSA Transformer with and without stochastic MSA sampling / ensembling, ESM3, ESM2, SaProt, TranceptEVE, RSALOR, Vespa, ESCOTT, and GEMME, in addition to NOMELT as a standalone call^18,24,31,51–63^. Scores from each model were compared to the experimental measurements of each point mutation in each incubation condition – computing Spearman’s, NDCG, and AUROC of improvement vs deleterious, all on a per position basis. The scores were then averaged over positions as well as recomputed on an entire library basis. We also considered Top K recovery over the entire library defined as number of top scoring variants required to recover K target mutations

For training set design, the full library was embedded using the ESM2-3B max pooling over mutable (not fixed) residues and transformed into principal component space at 98% variance – the same feature set as used for the final Round 1 supervised model. A custom implementation of the Frank-Wolfe algorithm was developed to maximize: *log*(*det*(*X*^!^ *X*)) where *X* is the feature matrix in PC space of currently selected variants and previously tested ones. The algorithm iteratively selects sequences that maximally increased D-optimality by computing gradients with respect to the Fisher information matrix and choosing the sequence providing the largest increase in D-optimality. Starting from the initial set of experimentally tested sequences (191 variants) plus the top 12 sequences identified by greedy prediction-based selection, we selected an additional 200 in this way, see the code on Zenodo [https://zenodo.org/records/18894815].

### Supervised Machine Learning Methodologies

All supervised training, hyperopt, and evaluation was conducted using AIDE^64^. For the first round of supervised training protein sequences were encoded using the ESM2-3B model (esm2_t36_3B_UR50D). Model sizes down to 35M were tested but 3B produced significantly improved performance. Embeddings were extracted from the final layer and processed using three pooling strategies across variable positions: flattening (concatenating all position embeddings), mean pooling, and max pooling. Max pooling was selected as the optimal strategy based on cross-validation performance. An ElasticNet regression model was implemented within a scikit-learn pipeline consisting of: (1) StandardScaler for feature normalization, (2) PCA for dimensionality reduction, and (3) ElasticNet regressor for prediction. A randomized hyperparameter search over 2,000 parameter combinations was conducted using 10-fold random cross-validation. The parameter space explored included: ElasticNet alpha (L1/L2 regularization strength): reciprocal distribution from 0.01 to 1.0. ElasticNet l1_ratio (balance between L1 and L2 penalties): uniform distribution from 0 to 1. Performance was assessed using Spearman rank correlation as the primary metric. An MLP was also tested as a model head but was unable to produce anything predictive despite heavy regularization and 2,000 random hyperparameter optimizations.

In the second round, we repeated training on all ∼400 tested combinatorial variants with several key enhancements. Multiple embedding approaches were systematically compared: ESM2-3B with both max pooling and position-specific flattening, SaProt structure-aware embeddings using the wild-type AlphaFold2 structure, one-hot encoding, and MSA Transformer zero-shot likelihood scores as an additional feature *a la* Hsu et al.^17^ Two modeling approaches were evaluated through randomized hyperparameter optimization: linear models using ridge regression with L2 regularization and nonlinear models using a multi-layer perceptron (MLP) with early stopping. For the nonlinear approach, an ensemble strategy was implemented using BaggingRegressor with multiple MLP estimators to provide additional regularization as overfitting was observed in single models. Key hyperparameters optimized included: regularization strength (alpha: 10⁻³ to 10³), MLP architecture (1-3 hidden layers of 100 units), activation functions (ReLU, tanh), and ensemble size. Each of the two architectures paired with each embedding strategy was optimized over 50 random hyperparameter trials.

### 14C-Leucine Radiolabeling

To measure protein concentrations using radioactivity, a concentration of 10 µM ^14^C-leucine (Revvity NEC279E050UC) was added to the cell-free expression reaction and incubated at 30°C overnight, as has been described before^65^. In brief, after incubation, 6 µL of the total cell-free expression reaction was treated with 6 µL 0.5 N KOH and incubated at 37 °C for 20 minutes. The remaining cell-free expression reaction was centrifuged at 14,000 g for 10 minutes and 6 µL of the soluble supernatant was treated with KOH as well and incubated at 37 °C for 20 minutes. 4 µL aliquots of the subsequent treated reactions were spotted onto two 96-well filtermats (Revvity 1450-421) in triplicate, including a zero-protein control, and dried under a heat lamp for 20 minutes. One of the two filtermats was then washed three times with 5% w/v trichloroacetic acid for 15 minutes at 4 °C, followed by a wash with 200 proof ethanol also at 4 °C. The washed filtermats were dried overnight. After the filtermats were dried, both the washed and unwashed filtermats were heated to 70°C using a heat plate and scintillation wax (Revvity 1450-441) was melted on top. The filtermats were then loaded into cassettes and radioactivity was then measured with a Revvity MicroBeta2 (MicroBeta2 Windows Workstation version 6.0.0.0).

### Size Exclusion Chromatography

Analytical size exclusion chromatography (SEC) was performed using an NGC Chromatography System (Bio-Rad, USA) equipped with a Superdex 200 Increase 10/300 GL column (Cytiva, USA) all equilibrated w/ storage buffer (90 mM Tris-HCl, 150 mM NaCl pH=8) at 4 °C. The column was calibrated using a low-molecular-weight marker kit (Cytiva, USA) and the void volume (V0) was determined using Blue Dextran 2000. All standards and samples were run at a flow rate of 0.5 mL/min and relative absorption was monitored at 280 nm over the retention volume.

### Circular Dichroism Spectroscopy

CD spectra were recorded on a JASCO CD spectrophotometer using a 1-mm pathlength Hellma cuvette. Protein samples were measured at a concentration of ∼0.5 mg ml in 90 mM Tris-HCl, 150 mM NaCl pH=8. Temperature melts were performed from 20 to 95 °C, and CD signals were acquired at 222 nm in 1 °C increments per minute with 10 s equilibration and 1 s digital integration. Wavelength scans (200–260 nm) were acquired at 20°C and 95 °C. Melting temperatures were determined as the inflection point of the slope change in the data.

## Code availability

Notebooks and data used to conduct all training, hyperparameter optimization, and filtering of ProteinMPNN libraries are available on Zenodo [https://zenodo.org/records/18894815]. The code for ProteinGym, used to compute zero-shot scores on the SSM library is available at [https://github.com/OATML-Markslab/ProteinGym]. The notebook used to benchmark using those scores is on Zenodo [https://zenodo.org/records/18894815].

## Data availability

All data presented in the manuscript are available in either the source data or uploaded to the Zenodo of the article (https://zenodo.org/records/18894815). Source data is provided within the paper.

## Acknowledgements

M.C.J., G.B., and A.S.K. acknowledge support from the Department of Energy grant DE-SC0023278. M.C.J. acknowledges the Air Force Office for Scientific Research grant FA9550-23-1-0420, the Army Research Office grant W911NF-23-1-0334, and the Stanford Doerr School Accelerator for support this work and equipment used.

This work was authored in part by the National Laboratory of the Rockies, operated by Alliance for Energy Innovation, LLC, for the U.S. Department of Energy (DOE) under Contract No. DE-AC36-08GO28308. Funding to EK and GTB was provided by the U.S. DOE, Office of Science, Biological and Environmental Research Office. The views expressed in the article do not necessarily represent the views of the DOE or the U.S. Government. The U.S. Government retains a nonexclusive, paid-up, irrevocable, worldwide license to publish or reproduce the published form of this work, or allow others to do so, for U.S. Government purposes.

## Author Contributions

Conceptualization: J.T.L., E.K., I.M., G.M.L., M.C.J.

Methodology: J.T.L., E.K., I.M., K.S.Z.., P.N., B.A.S., A.S.K.

Investigation: J.T.L., E.K., I.M., K.S.Z., P.N., S.S., K.G.K., A.T., B.A.S., A.S.K. Software: J.T.L., E.K., P.N., K.G.K.

Funding acquisition: D.M., G.B., M.C.J. Supervision: A.S.K., D.M., G.B., M.C.J.

Writing: J.T.L., E.K., I.M., S.S., G.M.L., A.S.K., G.B., M.C.J.

